# The herpes simplex virus type I deamidase enhances propagation but is dispensable for retrograde axonal transport into the nervous system

**DOI:** 10.1101/704338

**Authors:** Austin M. Stults, Gregory A. Smith

**Author notes:** Corresponding author: Gregory A. Smith, Ph.D., Professor of Microbiology-Immunology, Northwestern University Feinberg School of Medicine, 303 E. Chicago Ave., Chicago, IL 60611.

## Abstract

Upon replication in mucosal epithelia and transmission to nerve endings, capsids of herpes simplex virus type I (HSV-1) travel retrograde within axons to peripheral ganglia where life-long latent infections are established. A capsid-bound tegument protein, pUL37, is an essential effector of retrograde axonal transport and also houses a deamidase activity that antagonizes innate immune signaling. In this report, we examined whether the deamidase of HSV-1 pUL37 contributes to the neuroinvasive retrograde axonal transport mechanism. We conclude that neuroinvasion is enhanced by the deamidase, but the critical contribution of pUL37 to retrograde axonal transport functions independently of this activity.

**IMPORTANCE:** Herpes simplex virus type 1 invades the nervous system by entering nerve endings and sustaining long-distance retrograde axonal transport to reach neuronal nuclei in ganglia of the peripheral nervous system. The incoming viral particle carries a deamidase activity on its surface that antagonizes antiviral responses. We examined the contribution of the deamidase to the hallmark neuroinvasive property of this virus.

## INTRODUCTION

Mammalian viruses of the alpha-herpesvirinae subfamily initially infect a mucosal epithelium and then transmit to innervating sensory and autonomic nerve terminals (1). Virus-mediated fusion into axon terminals results in the deposition of the capsid and tegument proteins into the cytosol, with the majority of tegument proteins dissociating from the capsid. However, at least three tegument proteins, pUL36, pUL37, and pUS3, remain capsid bound (2-5). The pUL36 tegument protein directly binds to the pUL25 component of the capsid surface (6-9), and tethers pUL37, pUS3, as well as the host dynein/dynactin microtubule motor, to the capsid (10-13). Each of the capsid-bound tegument proteins has a distinct enzymatic activity: pUL36 houses a deubiquitinase in its amino terminus (14, 15), pUL37 houses a deamidase in its carboxyl terminus (16), and pUS3 is a serine-threonine protein kinase (17). Of these enzymes, only the pUL36 deubiquitinase is reported to contribute to the neuroinvasive property of these viruses (18, 19). Nevertheless, pUL37 is a critical component of the neuroinvasive apparatus (20), with an amino-terminal region essential for the delivery of incoming capsids to the neural soma by sustaining dynein-based microtubule transport in axons (21, 22).

The herpes simplex virus type 1 (HSV-1) pUL37 deamidase antagonizes innate cytosolic sensors including retinoid-acid inducible gene-I (RIG-I) and cyclic GMP-AMP synthase (cGAS) and is an important virulence determinant following peritoneal injection into mice (16, 23). In this report, we examine whether the deamidase specifically promotes HSV-1 invasion of the nervous system.

## RESULTS

### Confirmation of attenuated interferon suppression during infection with HSV-1 encoding a mutated deamidase

Zhao et al. previously identified two cysteines in HSV-1 pUL37, C819 and C850, as critical for catalytic deamidation of RIG-I *in vitro*, with C819 serving as the catalytic site (16). In the current study, we intended to mutate the catalytic site in pseudorabies virus (PRV) and HSV-1, but we noted that neuroinvasive herpesviruses within the varicellovirus genus of the alpha-herpesvirinae subfamily lack the catalytic cysteine (Fig. 1). Therefore, a cysteine-to-serine change was introduced at C819 in HSV-1 strain F that mimicked the design of the previously characterized catalytic mutant (16). A second HSV-1 mutant was produced encoding C850S. Neither mutant was impaired for pUL37 expression during infection (Fig. 2A). The C819S catalytic mutant triggered 3-fold increased interferon beta expression relative to the wild type upon infection of normal human dermal fibroblasts (NHDF), consistent with reports that the deamidase antagonizes interferon signaling (Fig. 2B) (16, 23). Repair of the C819S mutant (C819S>C) restored the wild-type phenotype.

**FIG 1:**
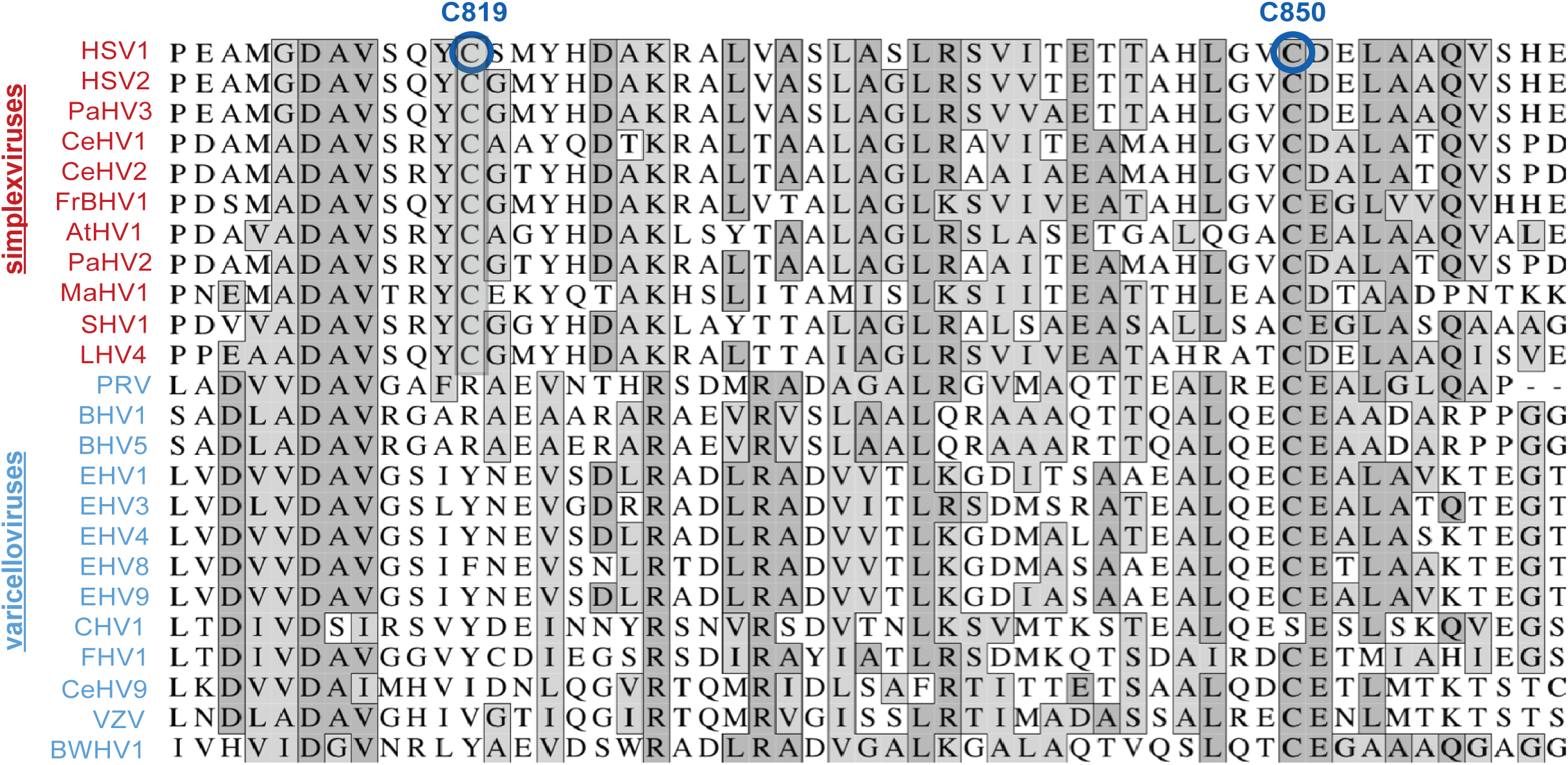
The pUL37 residues C819 and C850 are conserved within the simplexvirus genera. Alignment of the deamidase region of pUL37 across 24 members of the alpha-herpesvirinae subfamily spanning the simplexvirus and varicellovirus genera. The HSV-1 residues C819 and C850 are circled in blue.

**FIG 2:**
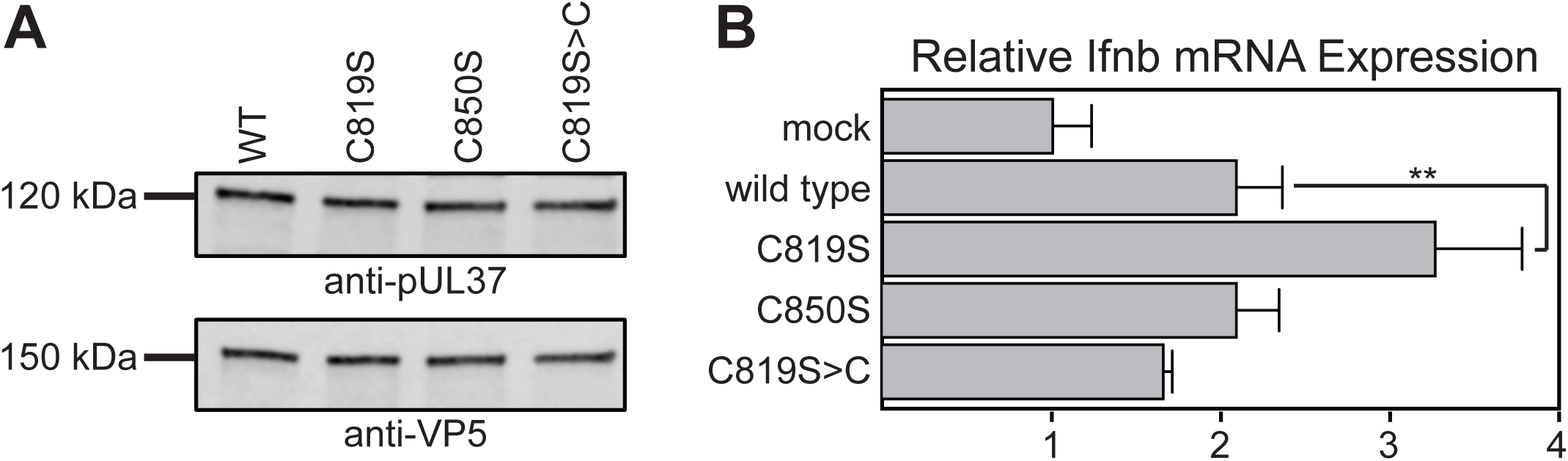
The catalytic residue of the pUL37 deamidase is required to antagonize interferon beta mRNA expression. (A) Western blot analysis of pUL37 expression in Vero cells at 18 hpi (MOI 10). (B) RT-qPCR analysis of interferon beta mRNA levels in NHDFs 5 hpi. Fold change is quantified relative to mock-infected cells. Error bars are s.d. (**, p < 0.01 based on ordinary one-way ANOVA followed by Dunnett’s multiple comparisons test).

Unexpectedly, HSV-1 encoding C850S was not defective for interferon suppression even though the residue was previously reported to support deamidase activity (16). The reason for this discrepancy was not clear, although we note that the previous study examined the C850S mutant during transient expression and did not examine the phenotype in the context of HSV-1 (16, 23).

### The pUL37 deamidase supports HSV-1 spread in culture

HSV-1 propagation kinetics were unaffected by either cysteine mutation (Fig. 3A); however, the C819S mutation reduced the spread of HSV-1 by 24% in primary fibroblasts and in Vero cells (Fig. 3B). To investigate the defect further, a recombinant of HSV-1 was produced encoding a CMV immediate-early promoter driving expression of the tdTomato fluorophore fused to a nuclear localization signal. Vero cells were infected at MOI 5 and harvested from 4-12 hpi to quantify the number of fluorescent cells by flow cytometry (Fig. 3C). A reduction in viral gene expression kinetics was observed for the C819S mutant virus, which reached 61.1 ± 0.6% of cells reporting at 12 hpi, compared to 89.7 ± 0.9% and 89.4 ± 1.4% for the wild-type and repair viruses, respectively. Because this result was not predicted by the single-step propagation results that were assessed at MOI 10 (Fig. 3A), Vero cells were infected at MOI 1, 5, and 10 and analyzed by flow cytometry at 8 and 24 hpi. The results indicated that the C819S defect was MOI dependent, with the greatest impact observed for MOI 1 and no defect at MOI 10 (Fig. 3C, right).

**FIG 3:**
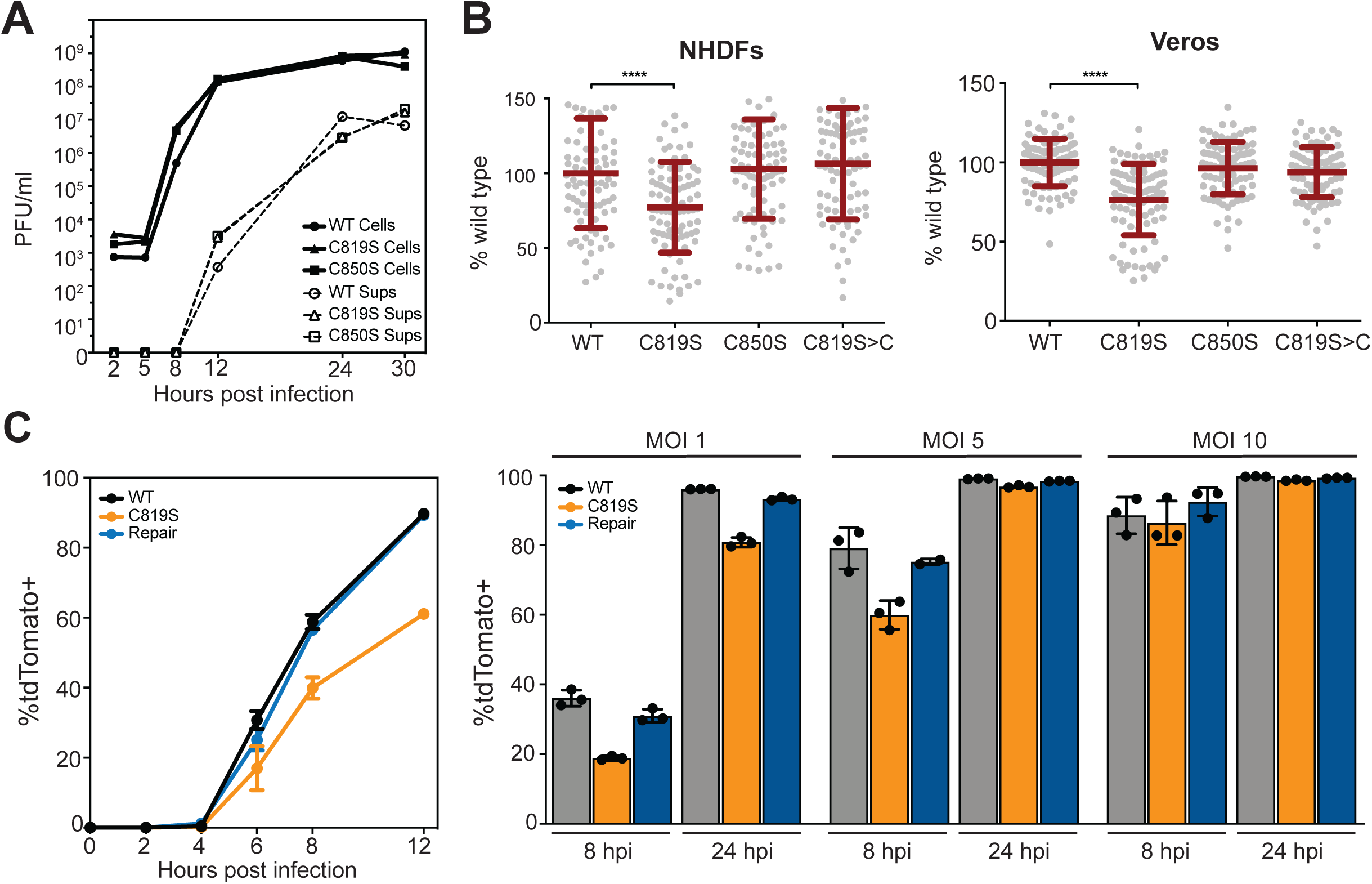
The pUL37 deamidase supports infection at low multiplicity and viral spread. (A) HSV-1 single-step propagation kinetics were determined by counting plaque-forming units harvested from Vero cells (Cells) and the corresponding supernatant (Sups) at the times indicated. (B) HSV-1 plaque sizes on NHDF cells at 32 hpi (left) and Vero cells at 72 hpi (right) were compiled across three independent experiments and plotted as a percentage of wild type. (C) Vero cells infected with HSV-1 encoding a tdTomato-NLS reporter at MOI 5 (left) or at the designated MOI (right) were harvested at the indicated times and scored for red fluorescence by flow cytometry. Error bars are s.d. (****, p < 0.0001 based on ordinary one-way ANOVA followed by Dunnett’s multiple comparisons test).

### The pUL37 deamidase supports HSV-1 propagation in the mouse cornea and neuroinvasion

Mice were ocularly infected with the wild-type and C819S viruses following corneal scarification to monitor propagation in the mucosa (tear film) and invasion of the peripheral nervous system (trigeminal ganglia; TG). Sampling of tear films by eye swab demonstrated that the C819S mutant and repair both expanded in the mucosa during the first 20 hpi and then retracted. However, the mutant expanded more slowly and to a lesser degree than the repair virus and retracted faster (Fig. 4A). The reduced propagation in the corneal mucosa correlated to decreased invasion of the TG (Fig. 4B). The wild-type virus consistently invaded the TG from the cornea, whereas invasion by the C819S mutant was stochastic. Several animals infected with C819S lacked detectable plaque-forming units in the TG but possessed viral DNA, consistent with the establishment of a dormant infection. The C819S repaired virus (C819S>C) and the C850S mutant virus were indistinguishable from wild type (Fig. 4C). To determine if the animals producing wild-type yields of the C819S mutant in the TG were the result of spontaneous reversion during infection, the plaque diameter of one such recovered virus was measured and found to be consistent with the C819S phenotype (Fig. 4D, left). This isolate was also confirmed to encode the C819S mutation (Fig. 4D, right).

**FIG 4:**
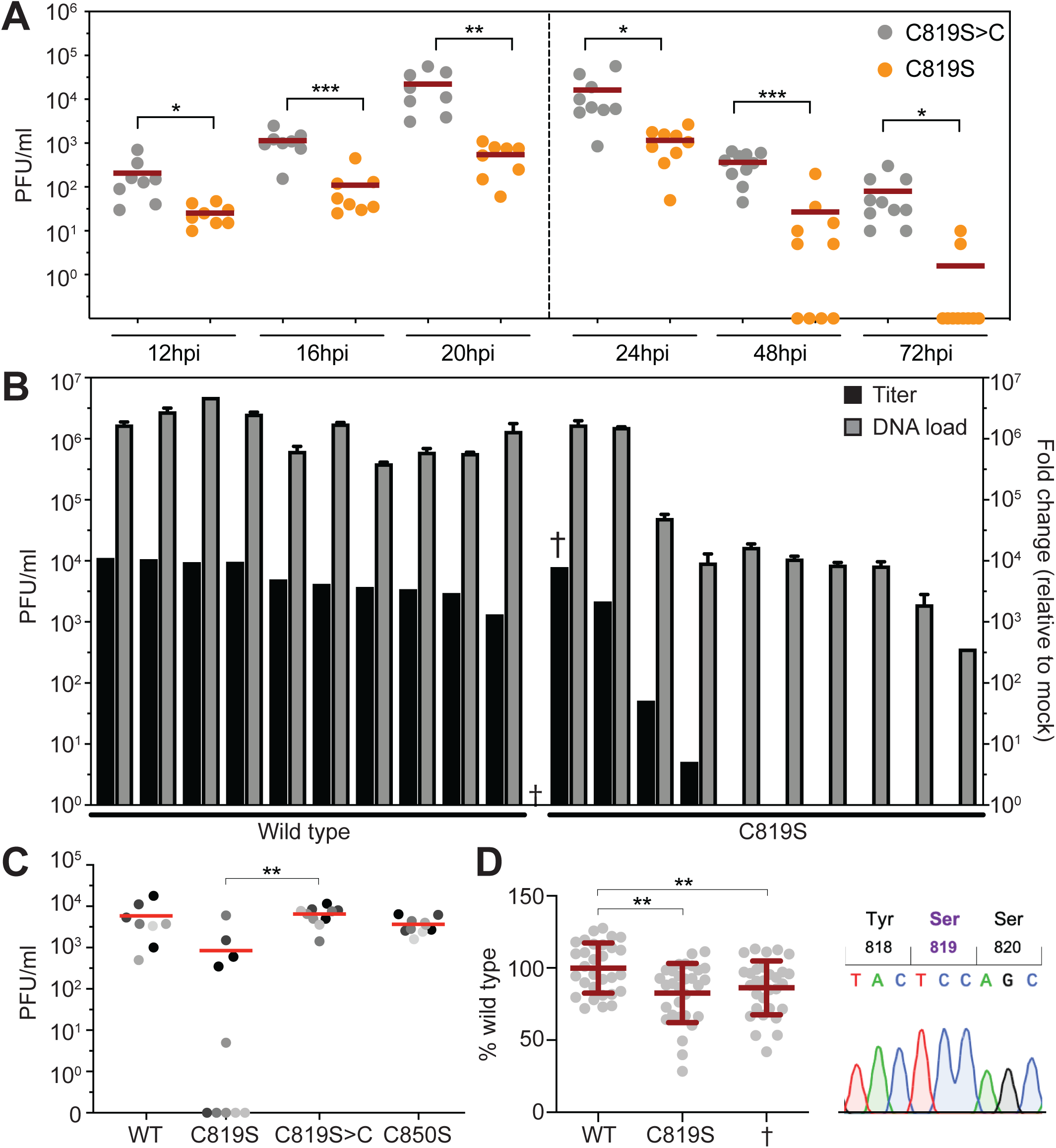
Mutation of pUL37 C819 delays invasion of the trigeminal ganglia upon ocular inoculation of mice. Mice were infected with HSV-1 encoding the pUL25/mCherry capsid fusion on both eyes following dual corneal scarification. (A) Tear film was sampled by swabbing each eye at the indicated times post infection. The data is a compilation of two experiments: four animals were used for the 12-20 hpi samples, and five animals were used for the 24-72 hpi samples. (B) Infectious HSV-1 (black bars) and HSV-1 DNA (grey bars) were recovered from individual TGs harvested and homogenized at 4 dpi. HSV-1 DNA levels were measured by qPCR and expressed as a fold change relative to mock-infected. Samples from each set of infections are presented in order of the titer recovered. (C) Titers were determined from TGs of additional mice. The mean titer of each virus is indicated by a red bar (10 TGs per virus from 5 mice; pairs of TGs from same animal are shaded equivalently). (D) Recovered C819S virus from the TG indicated in panel B (†) was examined for plaque diameter (left) and by sequence analysis (right). (*, p < 0.05; **, p < 0.01; ***, p < 0.001; based on two-tailed unpaired t test).

### Retrograde axonal transport is not dependent on the pUL37 deamidase

To test whether mutation of the deamidase had an effect on retrograde axonal transport, capsid transport dynamics were recorded within axons of primary sensory neurons during the first hour post infection. Mutation of either C819 or C850 had no significant impact on the directionality of capsid trafficking in axons (Fig. 5A), the average number of stops and reversals displayed by individual capsids (Fig. 5B), or the velocities and lengths of continuous retrograde runs (Fig. 5C). The distribution of retrograde velocities of each virus was consistently Gaussian (R^2^ ≥ 0.99 for each) and retrograde travel distances were accurately fit as decaying exponentials (R^2^ ≥ 0.91 for each), which is consistent with the processive motion of wild-type HSV-1 (5). Furthermore, delivery of capsids to the nuclear rims of primary sensory neurons was not notably impacted by either cysteine mutation (Fig. 5D). Collectively, these results indicate that the deamidase indirectly promotes neuroinvasion by enhancing HSV-1 propagation in peripheral tissues but does not directly contribute to retrograde axonal transport and delivery to neural soma in sensory ganglia.

**FIG 5:**
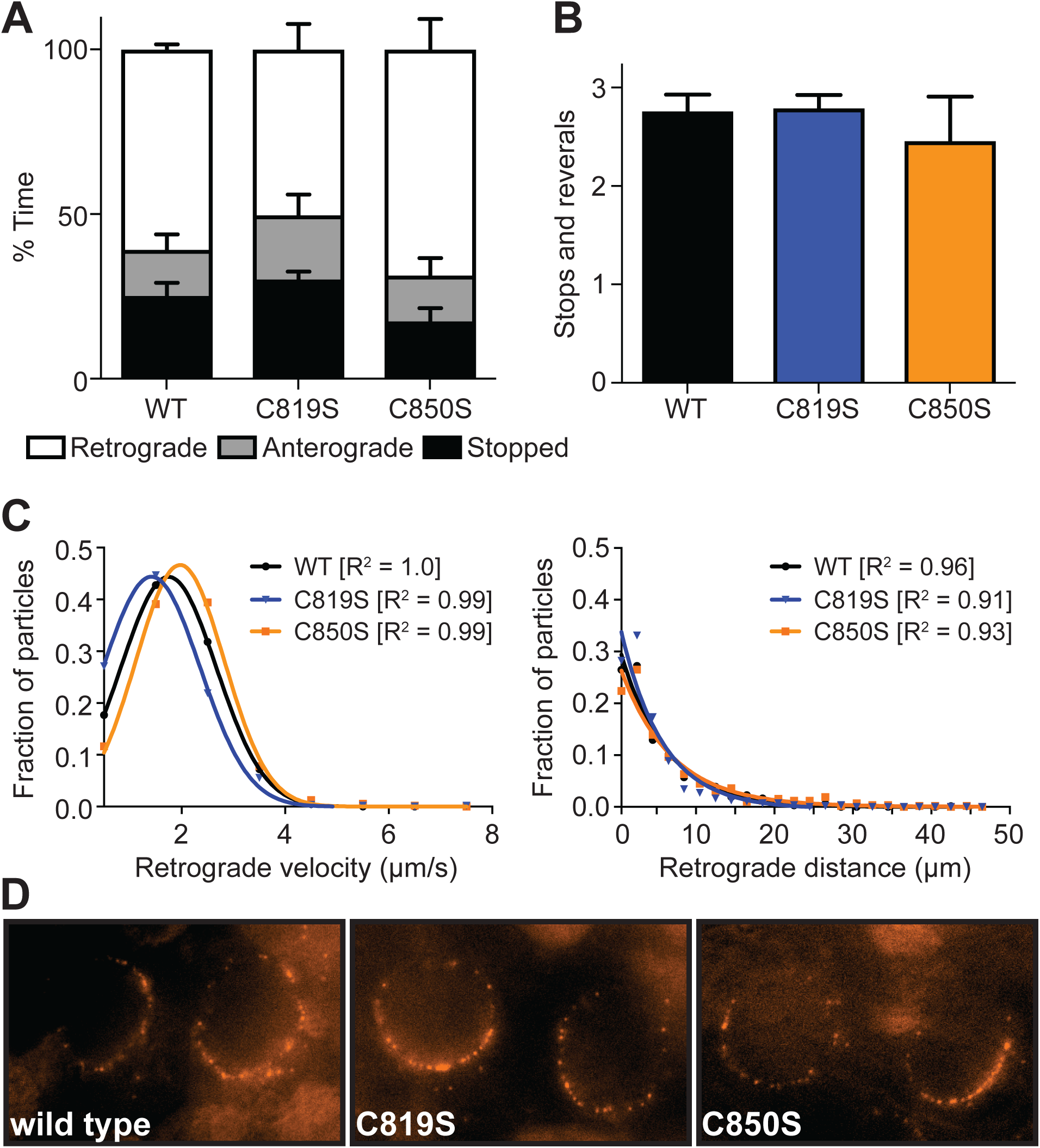
The pUL37 deamidase is dispensable for retrograde axonal transport. Primary sensory neurons were infected with HSV-1 encoding the pUL25/mCherry fusion and the indicated pUL37 allele, and transport dynamics in axons were monitored for the first hour post infection. (A) Fraction of time capsids moved in the retrograde direction, anterograde direction, or were stopped. (B) Average number of stops and reversals exhibited by capsids. (C) Distributions of forward run velocities and forward run distances of individual capsids in axons during the first hour post infection. (D) Representative images of capsids at nuclear rims of rat DRGs at 3-4 hpi. For panels A-D, more than thirty capsids were analyzed per experiment across three biological replicates.

## DISCUSSION

Viruses belonging to the simplexvirus and varicellovirus genres of the alpha-herpesvirinae subfamily are noted for their proficient neuroinvasive properties that result in the delivery of viral genomes to nuclei of peripheral ganglia neurons where latent infections are established and maintained. Upon initial exposure virus propagation within peripheral tissues, typically a mucosal epithelium, is required for robust transmission into the nervous system, presumably by increasing the number of viral particles encountering axon terminals at sites of innervation (24, 25). At least two viral proteins directly support the neuroinvasive mechanism. The pUL36 large tegument protein promotes transmission from epithelial tissues to nerve endings and subsequently tethers capsids delivered into the cytosol of axon terminals to the dynein/dynactin microtubule motor complex to drive retrograde axonal transport to sensory ganglia (12, 18). The pUL37 tegument protein sustains dynein-based microtubule transport by restraining opposing plus-end motion (22). Consistent with these roles, pUL36 and pUL37 remain attached to capsids upon entry into cells (2, 3, 5, 26), with pUL36 bound directly to the capsid surface (7, 9, 27, 28) and pUL37 bound to pUL36 (10, 11, 29) such that pUL36 tethers pUL37 to the capsid (13). Understanding how the pUL36 and pUL37 tegument proteins mediate neuroinvasion is foundational to developing vectors for neural gene delivery and vaccination.

The HSV-1 pUL37 tegument protein was recently found to house a deamidase activity that uses the cellular immune effectors, RIG-I and cGAS, as substrates to antagonize interferon-based antiviral responses (16, 23). The deamidase also supports HSV-1 dissemination into the brain following intraperitoneal challenge of mice, such that HSV-1 encoding a pUL37 C819S catalytic site mutation is incapable of transmitting to the brain from the peritoneum unless animals are knocked out for cGAS (23). This raises the question of whether the deamidase supports propagation in the periphery that indirectly enhances neuroinvasion, or if it also directly contributes to the neuroinvasive process. The latter possibility is compelling given that the pUL37 deamidase is anchored on the capsid surface during axonal trafficking (2, 5, 26).

In this report, we examined whether neuroinvasion is dependent on the deamidase using the mouse ocular model, which parallels the natural route of HSV-1 infection: inoculation of a mucosal site and subsequent retrograde axonal transport to the trigeminal ganglion of the peripheral nervous system. A pUL37 C819S mutant of HSV-1 strain F was produced to mimic the previously reported catalytic mutant and was confirmed to fail to antagonize interferon expression (16, 23). In mice, the C819S mutant displayed reduced propagation in the periphery that correlated to reduced neuroinvasion and production of recoverable virus from the trigeminal ganglion. The decreased propagation *in vivo* correlated to decreased propagation in culture, with the C819S mutant showing: (1) an increased dependence on multiplicity of infection to efficiently infect Vero cells, and (2) a decreased capacity to spread cell-to-cell in the plaque assay. Unexpectedly, the plaque deficit was noted in primary human fibroblasts as well as Vero cells, the latter of which are interferon deficient (30-32). The plaque results in Vero cells indicates that the pUL37 deamidase does more than antagonize interferon responses, or alternatively that RIG-I and cGAS trigger antiviral activities independently of interferon production.

Despite the propagation defects in culture and in mice, the C819S mutant was competent to invade the peripheral nervous system, and the capacity of the mutant to promote retrograde axonal transport in primary cultured sensory neurons was unimpaired. These findings raise the question of why the deamidase is housed in a capsid-bound tegument protein that is a critical effector of the transport mechanism. While only speculation can be offered on this point, it is noteworthy that the catalytic cysteine is absent from the neuroinvasive herpesviruses belonging to the varicellovirus genus (Fig. 1), suggesting that the deamidase may be absent from varicelloviruses. Nevertheless, representative members of these two neuroinvasive genera (HSV-1, simplexvirus; PRV, varicellovirus) engage in retrograde axonal transport dynamics that are indistinguishable from one another, which could indicate that the deamidase is a recent evolutionary adaptation of the simplexviruses and not a fundamental component of the neuroinvasive apparatus (5). In this regard HSV-1 also encodes an extended carboxyl terminus on pUL37 that modulates TRAF6 signaling (33), which is also absent from PRV. Together, these results demonstrate a role for the pUL37 deamidase in HSV-1 propagation and spread that is not directly required for retrograde axonal transport and invasion of the peripheral nervous system.

## MATERIALS AND METHODS

### Sequences and alignment

Predicted amino acid sequences of pUL37 homologs were aligned using the ClustalW alignment tool in MacVector. GenBank accession numbers used were: GU734771 (herpes simplex virus type 1; HSV1), NC_001798 (herpes simplex virus type 2; HSV2), YP_009011024 (panine alphaherpesvirus 3; PaHV3), NP_851897 (cercopithecine alphaherpesvirus 1; CeHV1), YP_164480 (cercopithecine alphaherpesvirus 2; CeHV2), BAP00716 (fruit bat alphaherpesvirus 1; FrBHV1), YP_009361900 (ateline alphaherpesvirus 1; AtHV1), YP_443884 (papiine alphaherpesvirus 2; PaHV2), YP_009227270 (macropodid alphaherpesvirus 1; MaHV1), YP_003933802 (saimiriine alphaherpesvirus 1; SHV1), YP_009230167 (leporid alphaherpesvirus 4; LHV4), JF797217 (pseudorabies virus; PRV), AJ004801 (bovine herpesvirus 1; BHV1), NC_005261 (bovine herpesvirus 5; BHV5), YP_053068 (equid herpesvirus 1; EHV1), YP_009054926 (equid herpesvirus 3; EHV3), NP_045240 (equid herpesvirus 4; EHV4), YP_006273002 (equid herpesvirus 8; EHV8), YP_002333504 (equid herpesvirus 9; EHV9), YP_009252247 (Canid alphaherpesvirus 1; CHV1), ALJ85051 (felid herpesvirus 1; FHV1), NP_077436 (cercopithecine alphaherpesvirus 9; CeHV9), NC_001348 (varicella-zoster virus; VZV), ASW27069 (beluga whale alphaherpesvirus 1; BWHV1).

### Recombinant HSV-1 production

All HSV-1 was derived from an infectious clone of HSV-1 strain F (34). A variant encoding an immediate-early gene reporter, HSVF-GS3217, was produced to monitor viral gene expression as an indication of viral genome delivery to nuclei. To make HSVF-GS3217, an En Passant template plasmid, pEP-CMV>tdTomato-NLS-in>pA, was first produced by modifying a CMV-driven eGFP cassette from pEGFP-N1 (Clontech), such that the CMV immediate early promoter was partially duplicated with an iscei::kan cassette inserted between the duplicated sequences. The entire merodiploid expression cassette was PCR amplified and the resulting linear dsDNA product was transformed into GS1783 bacteria harboring the full-length HSV-1 strain F infectious clone (35). Lambda red recombination was used to insert the product into the HSV-1 US5 gene (encodes the gJ glycoprotein), and the kan cassette was subsequently removed by a second round of lambda red recombination following digestion with the ISceI homing enzyme. Missense mutations (Table 1) were also introduced into the infectious clones by En Passant mutagenesis using primers listed in Table 2.

**TABLE 1:** Recombinant viruses

| Strain | Fluorescent reporter | Mutant | Titer (PFU/ml) |
| --- | --- | --- | --- |
| HSVF-GS4553<sup>a</sup> | pUL25/mCherry | - | 8.55 x 10<sup>7</sup> |
| HSVF-GS6801 | pUL25/mCherry | pUL37 C819S | 9.17 x 10<sup>7</sup> |
| HSVF-GS6769 | pUL25/mCherry | pUL37 C850S | 9.33 x 10<sup>7</sup> |
| HSVF-GS7016 | pUL25/mCherry | pUL37 C819S>C | 1.15 x 10<sup>8</sup> |
| HSVF-GS3217 | gJ::CMV>NLS-tdTomato>pA | - | 1.20 x 10<sup>8</sup> |
| HSVF-GS6958 | gJ::CMV>NLS-tdTomato>pA | pUL37 C819S | 1.73 x 10<sup>8</sup> |
| HSVF-GS6959 | gJ::CMV>NLS-tdTomato>pA | pUL37 C850S | 1.80 x 10<sup>8</sup> |
| HSVF-GS7065 | gJ::CMV>NLS-tdTomato>pA | pUL37 C819S>C | 1.23 x 10<sup>8</sup> |

<sup>a</sup> Previously published [Huffmaster 2015].

**TABLE 2:**
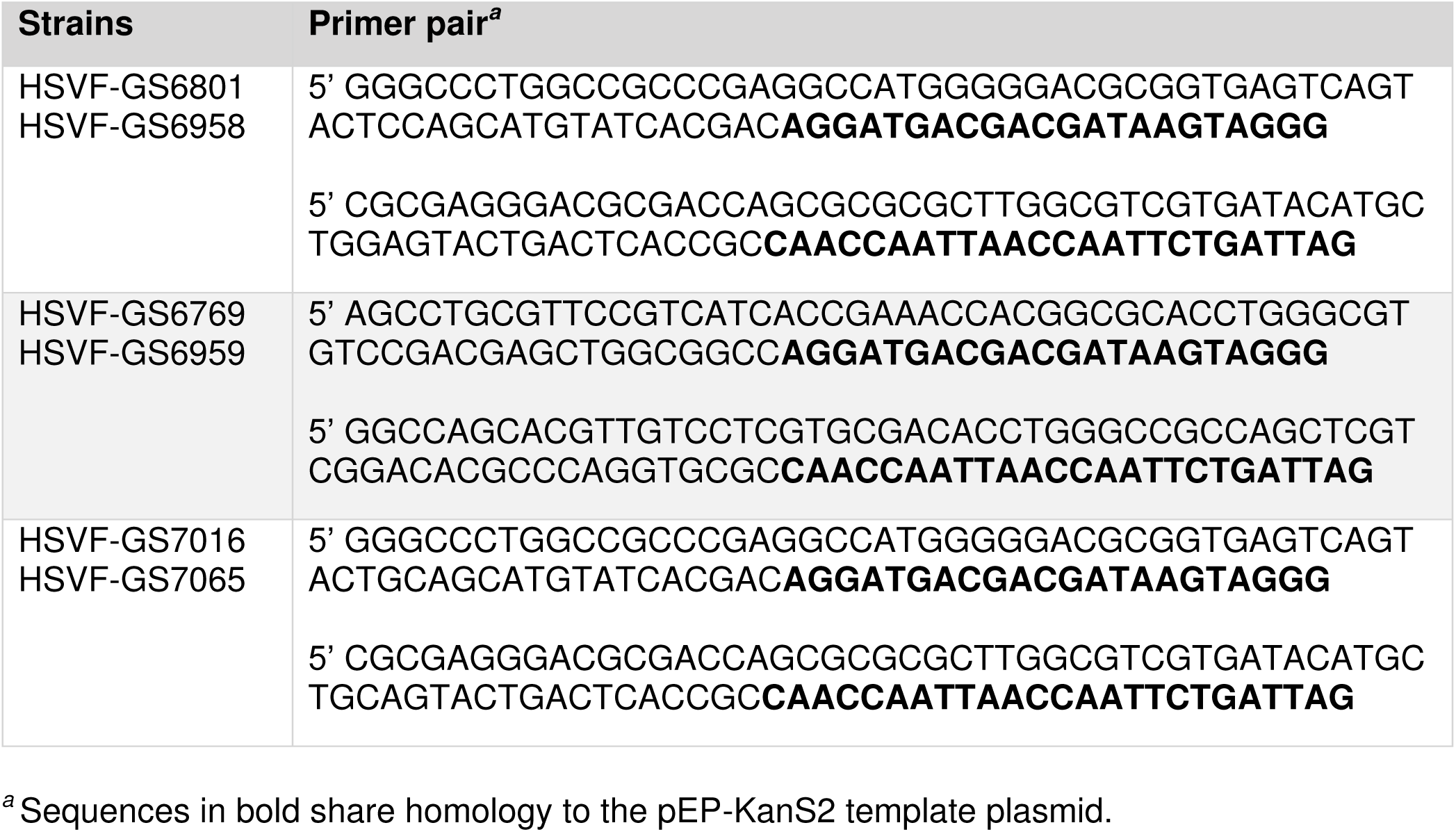
Primers used for En Passant mutagenesis

### Cell lines and HSV-1 propagation

Vero (African green monkey kidney epithelial, ATCC), Vero-CRE cells expressing Cre recombinase, and PK15 (pig kidney epithelial, ATCC) cells were grown in DMEM (Dulbecco’s Modified Eagle Medium, Invitrogen) supplemented with 10% BGS (bovine growth serum, RMBI). Normal human dermal fibroblasts (NHDFs) were generously provided by Derek Walsh and grown in DMEM supplemented with 10% FBS (fetal bovine serum, Gemini Bioproducts). Cells were tested regularly for mycobacterium contamination using the PlasmoTest kit (Invivogen) and authenticated by the source. BGS levels were reduced to 2% during and after infection. HSV-1 strains were produced by electroporation of infectious clones into Vero cells using an ECM630 electroporation system (BTX Instrument Division, Harvard Apparatus). Cells were pulsed once with the following settings: 220V, 950 µF, 0Ω. Serum levels were reduced to 2% BGS approximately 12 h after electroporation. Virus was harvested at a time at which 100% cells displayed pronounced cytopathic effect (CPE) (typically 5 days post electroporation). Initial viral harvests were subsequently passaged through Vero-Cre cells to excise the bacterial artificial chromosome vector from the viral genome (36).

### Single-step growth curve and plaque assays

Vero cells seeded in 6-well plates were infected at a multiplicity of infection (MOI) of 10. After 1 h, unabsorbed virus was inactivated with 1 ml of citrate buffer (pH 3.0), and cells were washed and incubated in 2 ml of DMEM supplemented with 2% BGS at 37°C, 5% CO_2_. At 2, 5, 8, 12, 24, and 30 hours post infection, HSV-1 was harvested from Vero cells and supernatants. Titers were determined by plaque assay on Vero cells overlaid with 2 ml methocel media (DMEM supplemented with 2% BGS and 10mg/ml methyl cellulose) and allowed to expand for five days. Images of at least 30 isolated plaques from each infection were acquired with a Nikon Eclipse TE2000-U inverted microscope fitted with a 0.30 numerical aperture (NA) 4 x objective and RFP filter set. To determine the plaque diameter, the average of two orthogonal diameter measurements was calculated for each plaque using ImageJ software. Plaque diameters were expressed as a percentage of the diameter of wild-type HSV-1, which was always measured in parallel. Data sets were plotted using GraphPad Prism 7 (GraphPad Software Inc).

### Primary neuronal culture

Dorsal root ganglia (DRG) from embryonic chicks (E8-E10) were cultured on poly-DL-ornithine- and laminin-treated coverslips in 2 ml of F12 media (Invitrogen) containing nutrient mix: 0.08 g/ml bovine serum albumin fraction V powder (VWR), 0.4 mg/ml crystalline bovine pancreas insulin (Sigma-Aldrich), 0.4 µg/ml sodium selenite (VWR), 4 µg/ml avian transferrin (Intercell Technology) and 5 ng/ml nerve growth factor (NGF; Sigma-Aldrich). DRGs from embryonic rats (E18) (Neuromics) were cultured as described above and supplemented with human holo-transferrin (Sigma-Aldrich). A single explant was cultured on each cover slip for 2 to 3 days and infected with 5 × 10^7^ PFU/ml of virus for five minutes. Cover slips were subsequently mounted to a glass cover slide and time-lapse imaging of mCherry emissions was achieved by automated sequential capture using 150 ms exposures between 0.5-1.0 hpi for retrograde transport analysis, and 3.0-4.0 hpi for nuclear rim formation.

### Fluorescence microscopy and image analysis

Virus transport dynamics were monitored in primary DRG explants. Explants were infected in 2 ml of F12 media with 5 × 10^7^ PFU/ml of HSV-1 (WT and mutants) from 0.5-1.0 hpi. Time-lapse images were captured using an inverted wide-field Nikon Eclipse TE2000-U microscope fitted with a 60x/1.4 NA objective and a CascadeII:512 electron-multiplying charge-coupled device (EM-CCD; Photometrics). The microscope was housed in a 37° environmental box (In Vivo Scientific). Moving particles were detected by time-lapse fluorescence microscopy in the red-fluorescence channel at 10 frames per second (continued 100 ms exposures) for 150 frames. Particle trajectories were traced in the 150 frame time-lapse image stacks using a multiline tool with a width of 20 pixels and average background subtraction, and a kymograph was produced using the MetaMorph software package (Molecular Devices). The multi-line tool was again employed to trace kymograph paths, and the fraction of time that a particle was stopped, moving anterograde, or moving retrograde was calculated for each particle. Forward distance and forward velocity for each particle were also measured and filtered for distances >0.5 microns to control for random diffusion of virions. Graphs were created in GraphPad Prism 7.

### Ethics statement

All procedures confirmed to NIH guidelines for work with laboratory animals and were approved by the Institutional Animal Care and Use Committee of Northwestern University (IS00003334). Fertilized chicken eggs were obtained from Sunnyside, Inc. and tissue was harvested between embryonic day 8 and 10.

### *In vivo* methods

Intraocular HSV-1 infections of BALB/c mice (9 week old; Jackson Lab) were carried out in animals anesthetized with an intraperitoneal injection of ketamine (86.98 mg/kg) and xylazine (13.04 mg/kg) mixture. Each cornea was lightly abraded 10 times in a crosshatched pattern with a 25-gauge needle, and 1 × 10^6^ PFU of HSV-1 was administered to the cornea surface. Prior to infection, the virus stock was sonicated and centrifuged for 2 min at 300 x g to remove cell debris. Tears containing shed virus were collected by proptosing each eye and swabbing with a damp cotton applicator three times in a circular pattern around the eye. Two independent experiments were performed to collect virus within the tear films of infected mice at 12, 16, and 20 hpi, and at 24, 48, and 72 hpi. At the indicated day post infection each trigeminal ganglion was removed and individually homogenized in 1 ml DMEM, sonicated, and stored at - 80°C. Titers of recovered HSV-1 from tissues were determined on Vero cells as described above.

### Preparation of cell lysates and Western blotting

Vero cells were seeded in a 6-well plate and infected at a MOI of 10. After 1 hour the inoculum was aspirated and replaced with DMEM + 2% BGS. After 18 hours the cells were washed once with ice cold 1X PBS and harvested in cold RIPA lysis buffer (50 mM Tris, pH 8, 150 mM NaCl, 1% NP-40, 0.5% sodium deoxycholate, 0.1% sodium dodecyl sulfate) supplemented with protease inhibitors (2.5 mM sodium fluoride, 1 mM sodium orthovanadate, 0.5 mM phenylmethylsulphonyl fluoride, 100 l of protease inhibitor cocktail [Sigma]). Lysates were rotated for 30 minutes, sonicated for three 1.5-s pulses, and rotated for an additional 30 min. Lysates were then spun at 13,000 X g for 20 min, and the supernatants were harvested. 2.5% of the supernatant from each lysate was mixed with 2X final sample buffer (62.5 mM Tris [pH 6.8], 2% SDS, 10% glycerol, 0.01% bromophenol blue) supplemented with 50 mM dithiothreitol (DTT) and loaded onto an acrylamide gel for SDS-PAGE. Blots were probed with an antibody raised against HSV-1 pUL37 (HA108, Virusys; diluted 1:3300) and an antibody raised against HSV-1 VP5 (diluted 1:2000; courtesy of Frank Jenkins).

### Genomic Sequencing

Viral DNA was isolated from stocks of infected Vero cells (titers ranging from 10^7^-10^8^ PFU/ml) using the PureLink Viral RNA/DNA kit (Invitrogen). Isolated viral DNA was used in a standard PCR reaction to amplify the genomic region encoding the deamidase, which was then purified using the Wizard Gel and PCR Clean-Up System (Promega) and submitted to a third party for sequencing.

### Quantitative PCR on mouse tissue

DNA was isolated from homogenized trigeminal ganglia using the DNeasy Blood and Tissue Kit (Qiagen). The final DNA concentration was diluted to 10 ng/μl. Each sample was run in triplicate using a 10 μl reaction volume consisting of: 5 μl of CyberGreen Mastermix (Roche), 0.5 μl forward primer (30 μM), 0.5 μl reverse primer (30 μM), 1.5 μl water, and 2.5 μl of DNA. Run settings were 95°C for 10 min, 50 cycles of 95°C for 15 sec, and 60°C for 30 sec. The forward and reverse primer sequences were: HSV-1 UL35 Fwd: GTCTTGGCCACCAATAACTC; HSV-1 UL35 Rev: GGGTAAACGTGTTGTTTGCG; mGAPDH Fwd: GATGGGTGTGAACCACGAG, and mGAPDH Rev: GTGATGGCATGGACTGTGG. Fold change was calculated using the 2^ΔΔCt^ method.

### Real-time quantitative PCR

RNA was isolated on ice from infected cells using the PureLink RNA Mini kit (Invitrogen). The final RNA concentration was diluted to 10 ng/μl and mixed with 10 μl of master mix to form cDNA. Run settings were 25°C for 10 minutes, 37°C for 2 hours, and 85°C for 5 minutes. cDNA was diluted 1:5 with RNAse-free water and used in a qPCR reaction as described above. The forward and reverse primers for each gene were previously published [Feng 2]: Human Ifnb Fwd: AGGACAGGATGAACTTTGAC; Human Ifnb Rev: TGATAGACATTAGCCAGGAG; Human B-Actin Fwd: CTGGCACCCAGCACAATG; Human B-actin Rev: GCCGATCCACACGGAGTACT. Fold change was calculated using the 2^ΔΔCt^ method.

### Viral gene expression

Vero cells were seeded to 100% confluency in six well plates and infected at a MOI 5 on ice for one hour to synchronize the infection of each virus. After one hour the cells were shifted to 37°C, 5% CO_2_. At the indicated times, cells were harvested, washed once with PBS, and fixed with 4% paraformaldehyde for 4-16 hours. Cells were then subject to fluorescence-activated cell sorting to quantify the percent of cells containing positive tdTomato signal. 20,000 cells were counted for each sample with each sample prepared in triplicate.

## ACKNOWLEDGEMENTS

We thank Sarah Antinone and Kevin Bohannon for their assistance in producing pEP-CMV>tdTomato-NLS-in>pA and HSVF-GS3217, and Derek Walsh for generously providing NHDF cells. Flow cytometry services were performed at the Robert H. Lurie Comprehensive Cancer Center Flow Cytometry Core, and sequencing services were performed at the Northwestern University Genomics Core Facility.

